# Arabidopsis annexin 5 controls plasma membrane properties in mature pollen grains

**DOI:** 10.1101/2021.03.02.433530

**Authors:** Małgorzata Lichocka, Magdalena Krzymowska, Magdalena Górecka, Jacek Hennig

**Affiliations:** Institute of Biochemistry and Biophysics, Polish Academy of Sciences, Pawinskiego 5a, 02-106 Warsaw, Poland

**Keywords:** annexin, pollen grains, pollen membrane, membrane repair, osmotic stress, ionic stress

## Abstract

In Arabidopsis, a dry stigma surface enables a gradual hydration of pollen grains by a controlled release of water. Occasionally the grains may be exposed to extreme precipitations that cause rapid water influx, swelling and eventually lead to pollen membrane (PM) rupture. In metazoans, calcium- and phospholipids-binding proteins, referred to as annexins participate in repair of the plasma membrane damages. It remains unclear, however, how this process is conducted in plants. Here, we examined whether the plant annexin 5 (ANN5), the most abundant member of the annexin family in pollen, is involved in the restoration of PM integrity. We analyzed a cellular dynamics of ANN5 in the pollen grains undergoing *in vitro* and *in vivo* hydration. We observed a transient ANN5 association to PM during the *in vitro* hydration that did not occur in the pollen grains being hydrated on the stigma. To simulate a rainfall, we performed spraying of the pollinated stigma with deionized water that induced ANN5 accumulation at PM. Similarly, calcium or magnesium application affected PM properties and induced ANN5 recruitment to PM. Our data suggest a model, in which ANN5 is involved in the maintenance of membrane integrity in pollen grains exposed to osmotic or ionic imbalances.

## Introduction

Pollen grains undergo extensive dehydration and enter a developmental arrest before their release from anthers. This state enables survival under harsh environmental conditions during pollination. In *Brassicaceace* the compatible pollen-pistil interaction requires several key steps for successful fertilization (Hiscock and Allen, 2008). After falling on the dry surface of stigma the pollen grains progressively absorb water. A recent study revealed that in Arabidopsis the intensity of water uptake by the pollen grains during hydration varies with time (Rozier et al., 2020). A rapid increase in the grain volume during the initial phase changes later on to less intensive. The controlled release of water by stigmatic papilla provides an effective mechanism enabling a selection of the most desirable pollen. When a pollen grain is rejected it fails to hydrate and consequently does not germinate (Chapman and Goring, 2010). By contrast, the proper hydration activates germination and formation of the pollen tube that extends through the tissues of a pistil toward the ovule. Reigning environmental conditions at the time of pollen dispersal may significantly affect successful pollination and fertilization. A heavy rain during the flowering is likely to reduce pollinators’ activity and evoke osmotic and mechanical stress in plants. It was shown that rain simulation by water spraying of Arabidopsis seedlings triggered a profound transcriptional reprogramming (Van Moerkercke et al., 2019). Moreover, the increased water absorption by pollen grains under conditions of high humidity significantly enhances a risk of pollen membrane disruption. To minimize this risk plants evolved membrane repair mechanisms. Synaptotagmin 1 (Syt 1) was the first component shown to play a role in this process (Schapire et al., 2008). More recent studies showed that Syt 1 acts at the plasma membrane-endoplasmic reticulum contact site conferring a mechanical stability to a plant cell (Perez-Sancho et al., 2015; Lee et al., 2019). Plasma membrane repair processes are better characterized in mammalian cells (Andrews and Corrotte, 2018) where annexins play an important role in restoring integrity of the plasma membrane after damage or injury (McNeil et al., 2006; Bouter et al., 2015; Demonbreun et al., 2019) and also upon osmotic shock (Creutz et al., 2012; Boye et al., 2017). The ability of annexins to self-assemble and form multimers at membrane damage site are prerequisites for membrane sealing. There has been a long-standing assumption that plant annexins may have similar properties (Schapire et al., 2009). Consistent with this model, an ectopic expression of some annexins reinforces plants tolerance to environmental cues improving their recovery after stress. A positive role in plant recovery after period of drought was shown for ANN1 and StANN1 in Arabidopsis and potato, respectively (Konopka-Postupolska et al., 2009; Szalonek et al., 2015); OsANN1 improved heat and drought tolerance in rice (Qiao et al., 2015); whereas overexpression of ZmANN33 and ZmANN35 promoted maize seedlings recovery after cold treatment and correlated with a reduced lipids peroxidation, a decreased levels of reactive oxygen species accompanied by an increased activity of peroxidases (He et al., 2019). However, direct contribution of plant annexins to membrane recovery has not been confirmed experimentally yet.

In Arabidopsis *ANN5* is predominately expressed in developing pollen grains and its essential for male gametophyte development (Zhu et al., 2014b; Lichocka et al., 2018). Interestingly, the homologous genes in *B. napus*, are mainly expressed in buds and newly formed pistils which might indicate species-specific function of the homologous annexins (He et al., 2020). Transcripts of other annexins, *ANN1, ANN2, ANN4*, were also detected during post-meiotic microspore development in Arabidopsis however only mRNAs of *ANN1* and *ANN5* are stored in the mature pollen grains (Honys and Twell, 2004; Wang et al., 2008). In contrast to *ANN1*, which is constitutively expressed in vegetative tissues and also during pollen development, the onset of *ANN5* expression occurs in the bicellular microspore and its transcripts accumulate until the pollen maturity (Honys and Twell, 2004; Zhu et al., 2014b). Previous studies showed that downregulation of *ANN5* level compromised pollen viability and vigor reflected by collapsing of the pollen grains and formation of the smaller grains with a reduced competitiveness compared to the wild-type grains (Zhu et al., 2014b; Lichocka et al., 2018). It was also shown that ANN5 *in vitro* binds to liposomes composed of phosphatidyl cholines and phosphatidyl serines in a calcium-dependent manner (Zhu et al., 2014a). However this observation has been not corroborated by experiments performed in living cells.

In this study, we show that ANN5 associates to membrane in pollen grains affected by osmotic or ionic stress. It binds adjacent to the damage sites supporting the model that plant annexins, similarly to their metazoan counterparts, are involved in membrane repair processes.

## Materials and Methods

### Plant material and growth conditions

The experiments were carried out on *Arabidopsis thaliana, Brasica napus, Nicotiana tabacum* and *Malus domestica* plants. *pLAT52:ANN5-YFP* and *pLAT52:ANN1-YFP* transformants were generated in Arabidopsis Col-0 background. *ANN5* RNAi-silenced lines 13 and 15 were previously generated in our lab (Lichocka et al., 2018). Arabidopsis plants were grown in Jiffy7 pots in controlled-environment chambers (Percival Scientific, Iowa, USA) at 22°C, ca. 55% relative humidity, under 8 h of light for 4 weeks followed by 16 h of light conditions, *B. napus* and *N. tabacum* plants were grown in soil under controlled environmental conditions (21 C, 16 h of light).

### Construction of expression vectors and transformation of Arabidopsis

*LAT52* promoter sequence (kindly provided from prof. Jörg Kudla) was PCR amplified using primers adding HindlII-XbaI restriction sites: forward 5’-AAGCTTCGACATACTCGACTC-3’ and reverse 5’-TCTAGAGTCCCCTTTAAATTGG-3’ and cloned into pJET1.2/blunt (Fermentas, St. Leon-Rot, Germany). The *LAT52* promoter sequence was subcloned into GWB441 bearing the full-length cDNA sequence of YFP (Yellow Fluorescence Protein). Coding sequence of *ANN5* was cloned as described previously (Lichocka et al., 2018) and LR recombined into modified GWB441(pLAT52). Coding sequence of *ANN1* was PCR amplified with primers adding SalI-Eco32I restriction sites: forward 5’-GTCGACATGGCGACTCTTAAGGTTTCTG-3’ and reverse 5’-GATATCCAGCATCATCTTCACCGAGAAG-3’ and cloned into pENTR1A vector compatible with the Gateway system. The resulting plasmid was LR recombined into modified GWB441(pL4T52). Wild-type Arabidopsis Col-0 plants were transformed using floral dipping method (Clough and Bent, 1998) and *Agrobacterium tumefaciens* strain GV3101 carrying the appropriate plasmid along with helper plasmid pMP90. Selection of T0 transformants was performed on solidified Murashige and Skoog (MS) medium (Duchefa, Amsterdam, The Netherlands) containing kanamycin (50 μg mL^-1^), 1.5% (w/v) sucrose and 1% (w/v) agar (Duchefa, Amsterdam, The Netherlands) in controlled-environment chambers at 22°C under 8 h of light. T1 plants (12 lines) were transferred from selection plates to Jiffy7 pots for cultivation. Further selection was performed on dry mature pollen grains that were spread on microscope slide and analyzed by fluorescence microscopy. We focused on YFP fluorescence intensity as a plant selection factor. Transgenic lines showing high number of transgenic pollen grains (ca. 90%) and high level of YFP fluorescence were selected for further propagation (five lines per genotype). Preliminary imaging of ANN5-YFP was performed on pollen grains from two T2 lines number 4 and 7 that revealed the same ANN5 distribution pattern. The results presented in this report were obtained for line number 4.

### *In vitro* pollen hydration and germination

To analyze ANN5 localization during pollen hydration dry pollen grains were placed between two cover glasses, 50 x 24 mm and 24 x 24 mm. Pollen specimen was mounted on inverted confocal microscope and time-lapse imaging was started before the addition of liquid medium. A small amount (30 μL) of 20% sucrose was added between glasses. Pollen grains hydrated rapidly increasing their volume. Enlarged grains formed an effective barrier that prevented further spread of the liquid between two cover glasses and consequently did not allow neighboring grains to hydrate. Pollen hydration was imaged for 2 min using Nikon C1 confocal microscope (Nikon Instruments B.V. Europe, Amsterdam, The Netherlands).

Freshly made pollen germination medium (20 mL) was composed of 20% (w/v) sucrose, 1.6 mM boric acid (H_3_BO_3_), 1.5 mM Ca(NO_3_)_2_, 1 mM MgSO_4_, 1 mM KNO_3_; pH 7.5 was adjusted with NaOH. Mature pollen grains were spread on chambered cover glasses followed by the addition of 200 μL liquid germination medium. FM4-64 was added to the pollen sample to a final concentration of 0.01 μg mL^-1^to counterstained pollen tube membrane. After 30 min incubation at room temperature pollen tubes were examined under confocal microscope.

### Pollination assay

Dry pollen grains overexpressing *ANN5-YFP* were applied over a small area on the rectangular cover glass (24 x 50 mm). To keep stigma alive for at least 30 min, a small block (3 × 4 × 2 mm) of a solid MS medium was placed on the cover glass near the area where pollen grains were applied. The emasculated pistil with the pedicel from flower bud stage 12 was mounted so that the pedicel was lying on the agar block whereas the stigma papillae on the cover glass in the area where pollen grains were applied. Confocal images were acquired every 1 min over a total of 25 min.

### Analysis of pollen membrane properties

Freshly prepared sucrose solutions (20% (w/v)) were supplemented with the individual components of the pollen germination medium to a final concentration of 1.5 mM Ca(NO_3_)_2_, 1 mM MgSO_4_, 1 mM KNO_3_ or 1.6 mM H_3_BO_3_. Next, mature pollen grains of the wild-type or overexpressing *ANN5-YFP* were spread over chambered cover glasses and subsequently of 200 μL of appropriate solution was added. Changes in PM permeability were monitored with the use of propidium iodide (PI) (Thermo Fisher Scientific, USA) that was added after 5 min incubation in appropriate solution to a final concentration of 0.25 μg mL^-1^. Change in the PM potential was monitored in wild-type pollen grains with a the use of DiOC2(3) (Thermo Fisher Scientific, USA) that accumulates in hyperpolarized membranes and DiBAC(4)3 (Thermo Fisher Scientific, USA) that enters depolarized cells (Suchodolski and Krasowska, 2019). The voltage sensitive dyes were added separately to the pollen grains after 5 min incubation in appropriate solution to a final concentration of 0.005 μg mL^-1^ for both dyes. The time-lapse imaging of pollen grains was started 5 min after the addition of the dye and continued for the next 30 min.

### Confocal laser scanning microscopy

Subcellular localization of ANN5-YFP fusion proteins in pollen grains was evaluated using a Nikon C1 confocal system built on TE2000E and equipped with a 40× PlanFluor and 60× Plan-Apochromat oil immersion objective (Nikon Instruments B.V. Europe, Amsterdam, The Netherlands). Fluorescence of YFP, DiBAC(4)3 and DiOC2(3) was excited with a Sapphire 488 nm laser (Coherent, Santa Clara, CA, USA) and observed using the 515/530 nm emission filter. FM4-64 and propidium iodide fluorescence were excited with a 543 nm HeNe laser and detected using the 610LP nm emission filter. Imaging of samples with two different fluorophores was performed in sequential mode to prevent signal bleed-trough. For publication, images were collected in single plane or z-stack mode at a 0.8 μm-focus interval. Z-stacks were rendered into 2D image in maximum intensity projection mode using ImageJ.

### Image processing and quantification

Confocal images were processed with the object separation procedure in ImageJ before quantification of the individual pollen grains area (https://imagej.net/Nuclei_Watershed_Separation). First, images were converted to 8-bit monochrome images. Next, Gaussian blurring was applied at the default settings. Further, pixel intensity tresholding was performed to separate of the objects from the background. We used black&white tresholding that resulted in white background and black objects. To separate individual pollen grains on tresholded images Watershed algorithm was applied. Segmented pollen grains were selected manually with Wand tracing tool and added to the ROI Manager tool. Next area of selected grains was measured [μm^2^]. Pollen grains were selected manually instead by Analyze Particle tool because only few grains per image were suitable for measurements. We rejected objects with not accurate confocal cross-sectioning of the grain area.

### Statistical analysis

Statistical analyses were conducted in R Studio software environment, version 1.2.5033, employing R software, version 3.6.3. (R Core Team, 2020). First, data was tested for the normality of distribution by Shapiro – Wilk test. Distribution was also validated by Q–Q (quantile-quantile) plots. The homogeneity of variances was evaluated by Levene’s test. If requirements for parametric tests were met, ANOVA (one-way or two way) was used. Otherwise, Kruskal – Wallis test performed. For post-hoc testing Tukey HSD, or Wilcoxon test were applied, respectively. Plots were generated using ggplot2 package (Wickham, 2016).

## Results

### Osmotically-driven assembly of ANN5 on pollen membrane

In this study we focused on a role of ANN5 in mature pollen during hydration and germination. In order to monitor its distribution we generated Arabidopsis lines expressing ANN5 fused to YFP under the control of *LAT52*, a pollen-specific promoter (Twell et al., 1990). It was shown that *in vitro* hydration of the dry pollen may induce a transient leakage of the pollen membrane (PM) due to a rapid membrane overstretching (Crowe et al., 1989). Since some metazoan annexins are involved in membrane repair, we wondered whether ANN5 of Arabidopsis could associate to PM exposed to overstretching. Sucrose solution (20%), which was shown to be appropriate for *in vitro* germination of Arabidopsis pollen (Zhu et al., 2014a), was used for pollen hydration. The pollen grains from freshly opened flowers, collected up to 48 h after onset of anthesis were placed between two cover glasses. Time-lapse imaging of ANN5 distribution was started before the addition of the sucrose solution and continued for 2 min. The pollen grains hydrated rapidly and attained their final volume within 40 seconds (Fig. 1A). We observed a rapid recruitment of ANN5 to PM during the late period of water uptake (Fig. 1B). This reaction was transient, and as soon as the pollen grains had reached an appropriate hydration level, ANN5 was progressively released from the membrane. In the fully hydrated pollen grains ANN5 was localized to the pollen cytoplasm and vegetative nucleus.

**Figure 1.**
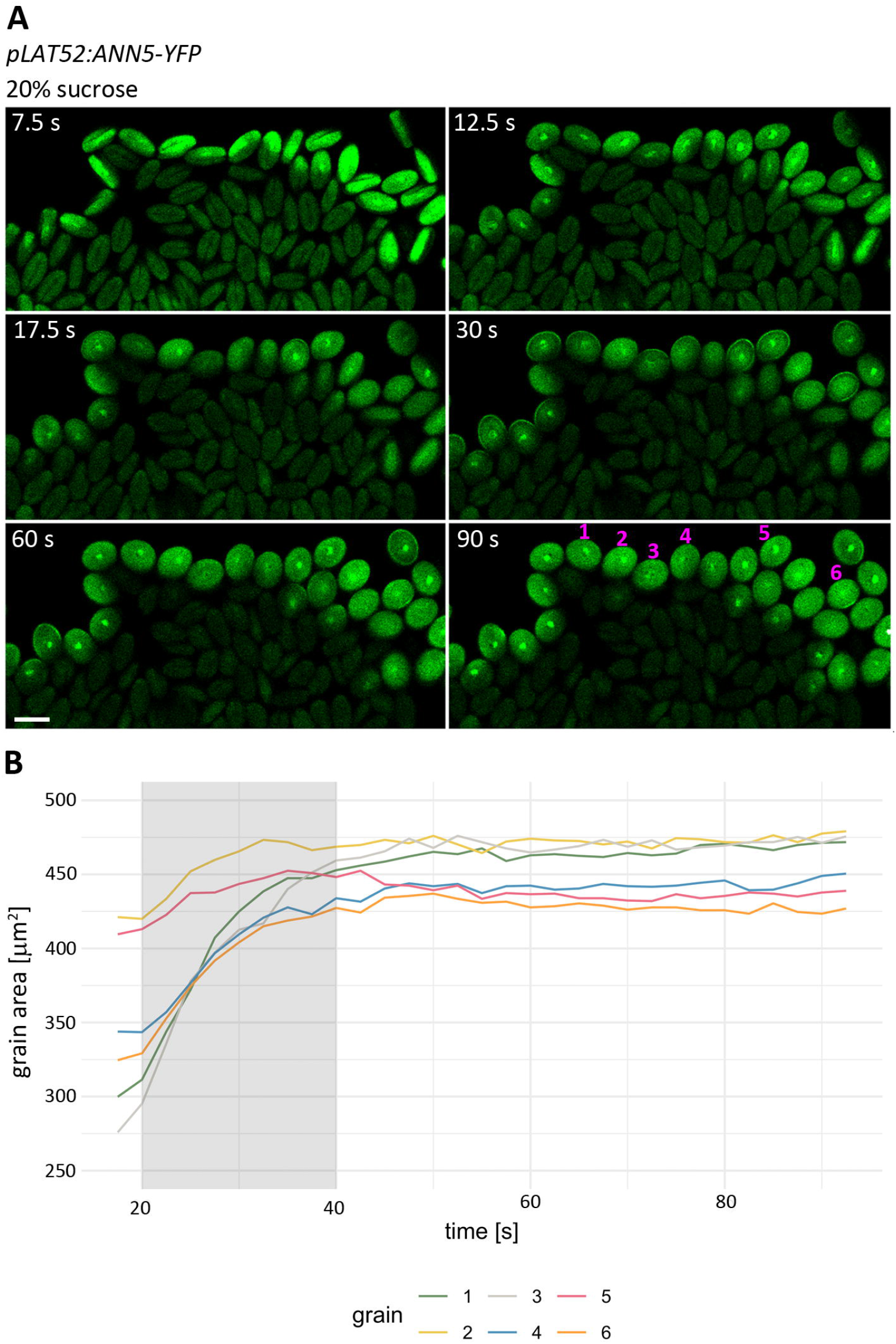
Transient association of ANN5 with PM in mature pollen grains during *in vitro* hydration. **(A)** Representative time-lapse images showing ANN5 distribution in pollen grains overexpressing *ANN5-YFP* during hydration in 20% sucrose. Confocal imaging was started before addition of sucrose solution, and the time [s] elapsed after adding of medium is indicated on each image. A scale bar = 20μm. (**B**) Plot of the change in pollen grains area [μm^2^] during *in vitro* hydration. Pollen grains marked with numbers (1-6) were selected for the measurement of the grains area on single time-lapse images. The gray shaded region indicates the duration of ANN5 attachment to PM [s].

Interestingly, a prolonged incubation of the pollen grains in the sucrose solution resulted in re-recruitment of ANN5 to PM in some grains (Fig. 2A, 2B). We set out to verify whether there is any correlation between binding of ANN5 to PM and loss of PM integrity. To this end, pollen grains were stained with propidium iodide (PI) that is a membrane impermeant dye, which enters cells upon the membrane damage. We found that binding of ANN5 to the PM occurred in the pollen grains with compromised integrity of PM, identified by positive PI staining. There were, however, grains in which ANN5 was associated with PM although PI influx was not detected suggesting that ANN5 binds to osmotically-stressed pollen membranes before a measurable loss of their integrity (Supplementary Fig. S1).

**Figure 2.**
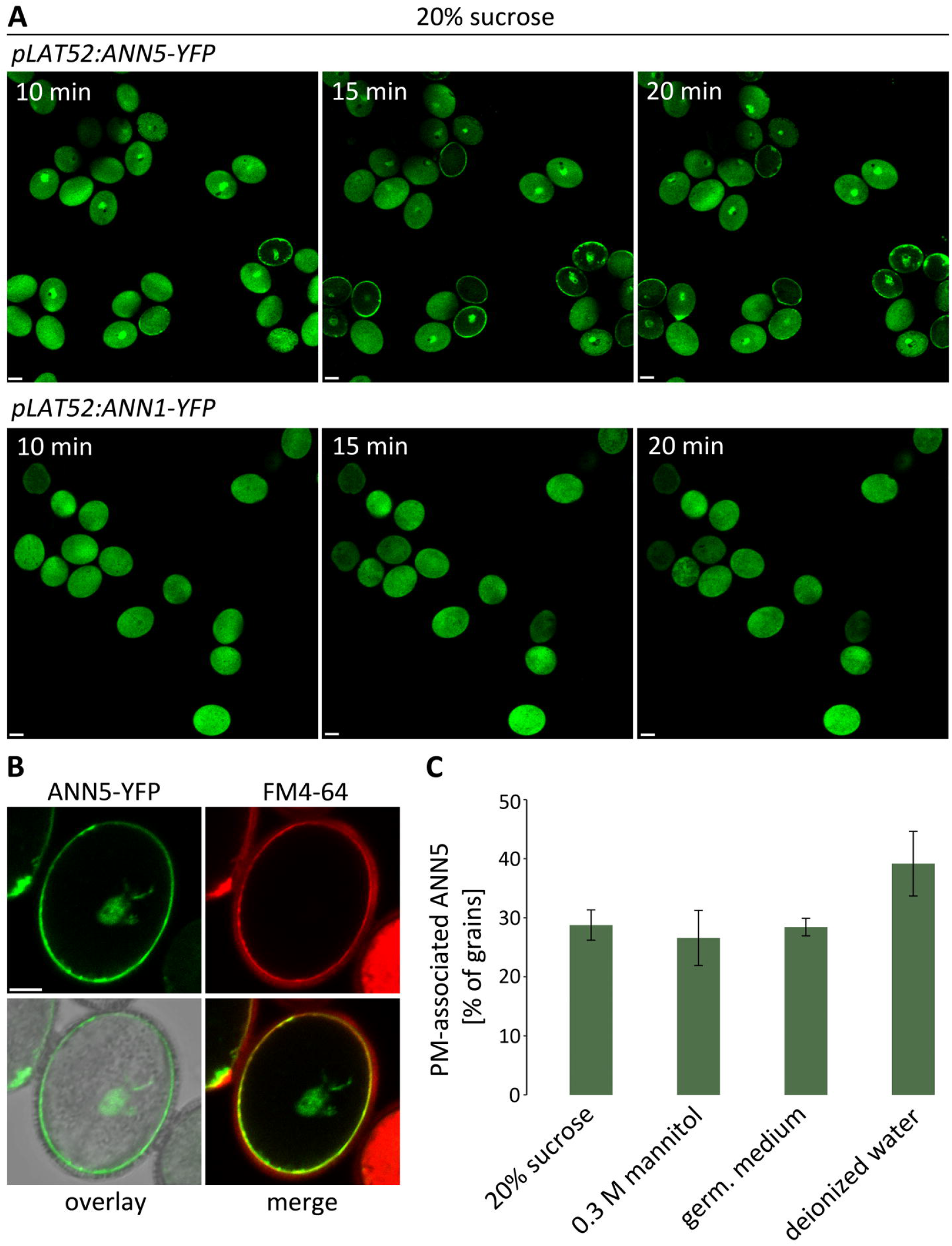
The cellular dynamics of ANN5 and ANN1 differ in response to osmotic stress. **(A)** Comparison of the subcellular localization of ANN5 and ANN1 in mature pollen grains during incubation in 20% sucrose. Representative images showing accumulation of ANN5 at pollen membrane (PM) that did not occur in pollen grains overexpressing *ANN1-YFP*. Scale bars = 10 μm. See also Supplementary Fig. S1. **(B)** Representative image showing colocalization of ANN5 with the lipophilic fluorescent dye FM4-64 at PM in osmotically stressed pollen grain. A scale bar = 5 μm. **(C)** Fraction of pollen grains [%] showing PM-associated ANN5 after 30 min incubation in deionized water, 300 mM mannitol solution, 20% sucrose and liquid pollen germination medium. The data were obtained in three independent experiments. n ≥ 400 pollen grains in each solution. Bars represent SD. No significant differences between treatments were detected.

To test the effect of other osmolytes on ANN5 distribution we performed a similar assay using mannitol solution at a concentration of 300 mM, that was shown to provide iso-osmotic environment for a lily (*Lilium longiflorum*) pollen and is commonly used for sampling of Arabidopsis pollen (Pertl et al., 2010; Lu, 2011). In this experiment we also applied a standard liquid germination medium that provides osmotic balance for pollen grains and deionized water to induce hypo-osmotic stress. Live-cell imaging of pollen grains was started following 30 min incubation in the appropriate solution and continued for the next 30 min. Standard pollen germination medium, 20% sucrose solution or 300 mM mannitol should promote pollen hydration and survival of the majority of pollen grains. In those grains ANN5 was localized to the pollen cytoplasm and vegetative nucleus. However, in a substantial pool (approximately 25%) of the pollen grains ANN5 displayed PM localization under such conditions (Fig. 2C). As expected, the number of PM-associated ANN5 grains incubated in deionized water was higher than in iso-osmotic solutions. However, the majority of the pollen grains survived and maintained cell integrity under hypo-osmotic stress. We hypothesize that some pollen grains have reduced ability to sustain adverse environmental condition. We assume that in those grains the osmotic equilibrium state was not achieved and in consequence a large cytoplasmic pool of ANN5 was shifted to the PM to protect the pollen grains from rupture.

Considering that transcripts of *ANN1*, a ubiquitously expressed annexin, were also detected in the mature pollen grains, we wondered whether ANN1 might display a similar cellular dynamics. To investigate this possibility we analyzed distribution of ANN1 in the mature pollen grains incubated in the sucrose solution for 20 min. ANN1 was localized to the pollen cytoplasm and its distribution remained unchanged over the experiment (Fig. 2A). This finding indicates that ANN1 and ANN5 play distinct roles in the mature pollen grains and only ANN5 may exert protective effects on PM. Therefore the ANN5 involvement in the in the restoration and maintenance of PM integrity was further analyzed in detail.

### Simulation of rainfall induces translocation of ANN5 to pollen membrane in pollen grains germinating on stigma

*Brassicaceae* possess a dry-type stigma that after pollination supplies water for pollen hydration and germination. To determine whether gradual hydration of pollen grains would have an impact on ANN5 distribution we performed live-cell imaging of pollen grains overexpressing *ANN5–YFP* after being placed on the stigma surface. To this aim we developed a sample preparation protocol for imaging of single pollen grain on papilla cell under an inverted confocal microscope (see Materials and Methods). Once the pistil was mounted on the cover glass live-cell imaging was started. The initial phase of the pollen hydration on the stigma was slower in comparison with the application of the sucrose solution. This resulted in the partial pollen grain hydration similarly as recently observed by Rozier et al. (2020). Consequently, PM was not exposed to overstretching or damage thus ANN5 remained cytoplasmic, while its pool localized to the vegetative nucleus (Fig 3A). We observed a local increase in YFP fluorescence intensity during the establishment of a germination site and in the elongating pollen tube that suggests also an ANN5 involvement in the pollen tube formation

**Figure 3.**
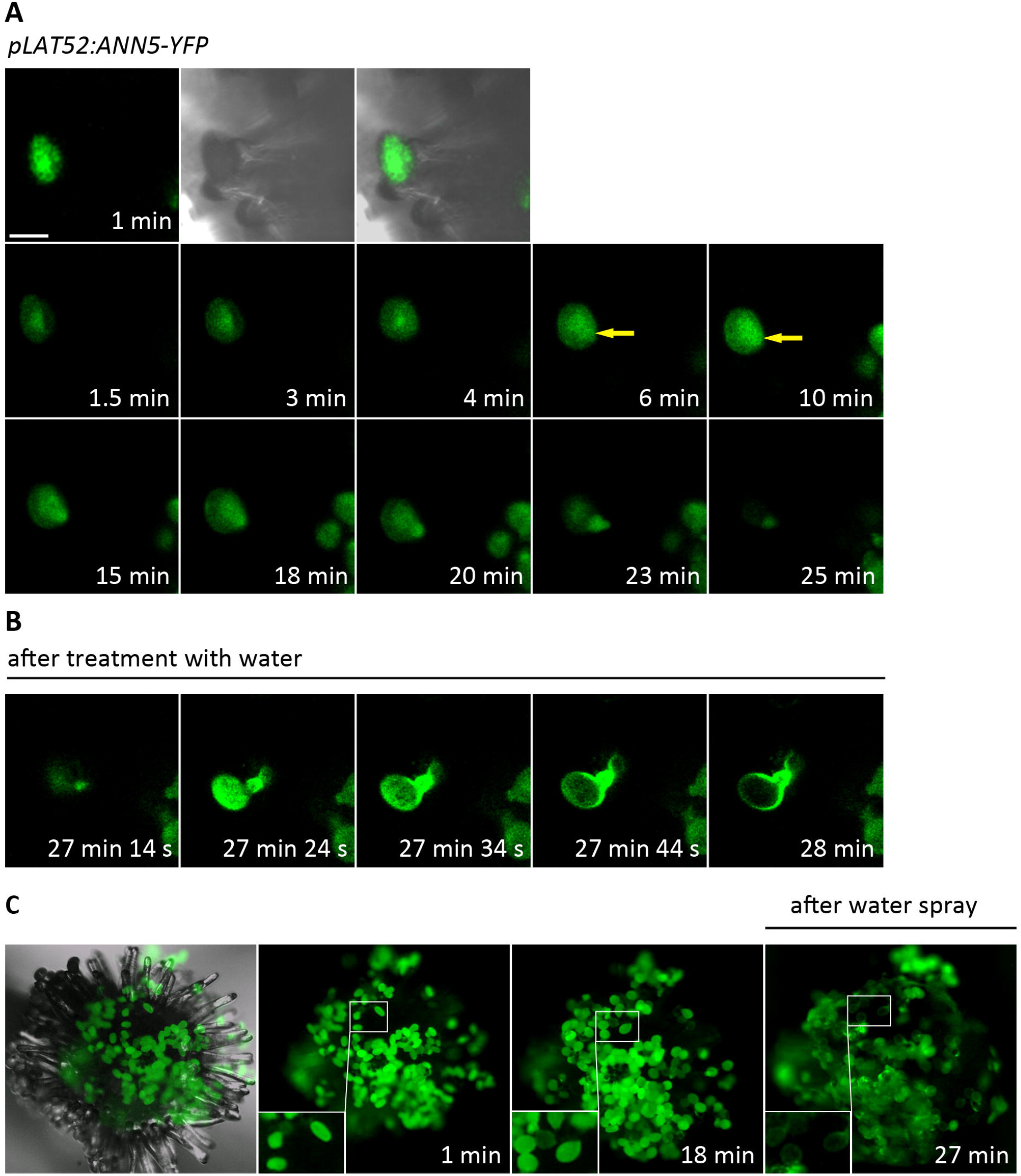
Recruitment of ANN5 to PM after treatment of pollinated stigma with deionized water. **(A)** Representative time-lapse images of single pollen grain overexpressing *ANN5-YFP* acquired during hydration on stigma. Yellow arrows indicate local accumulation of ANN5 corresponding to the pollen germination site. A scale bar = 20 μm. (**B**) Excerpts from a movie showing a translocation of ANN5 to PM after placing a drop (30 μl) of deionized water on pollinated stigma. See also Supplementary Video S1. (**C**) Fluorescent stereomicroscope images of stigma pollinated with pollen grains overexpressing *ANN5-YFP*. Images were acquired 1 and 18 min after hand-pollination of stigma. After 25 min pollinated stigma was sprayed three times with deionized water at the approximate distance of 25 cm. Image showing clumping of pollen grains and ANN5 translocation to PM was captured 2 min after treatment with water.

Under natural conditions the water balance can be suddenly changed by a rainfall. The excessive water uptake may induce bursting of pollen grains and pollen tubes, preventing successful fertilization. To simulate rain, we applied a drop (30 μL) of deionized water on the pollinated stigma that induced a rapid and massive translocation of ANN5 to PM (Fig. 3B, Supplementary Video S1). Similar results were obtained in other live-cell imaging experiment in which emasculated pistils were mounted on agar plate, manually pollinated and then sprayed with deionized water during pollen tube formation. Using stereo fluorescent microscope we got a more complete picture of the changes induced by application of water onto the pollinated stigma. We observed clumping of the pollen grains and papilla cells and translocation of ANN5 to PM in numerous pollen grains (Fig. 3C). Taken together, the data suggest that ANN5 may support the PM integrity during swelling of pollen grains induced by an excessive water uptake.

### ANN5 binds to osmotically disrupted pollen tube membrane

Considering that we observe recruitment of ANN5 to the pollen germination site and its accumulation in the pollen tubes we set out to perform more detail characterization of the ANN5 distribution in pollen tubes. To this end, we monitored pollen grains germination in a standard liquid pollen germination medium under inverted confocal microscope. Counterstaining with FM4-64 was performed for tracking the dynamics of membranes in the growing pollen tubes. After 30 min incubation at room temperature we found that none of the grains with ANN5 attached to PM germinated (Fig. 4A). In contrast, the grains where ANN5 was distributed in the pollen cytoplasm, formed pollen tubes. Along with the extension of the pollen tube we observed an increased fluorescence intensity at the tube apex due to the accumulation of ANN5 (Fig. 4B). ANN5 co-localized with vesicle-rich areas in this region, marked by FM4-64 staining. Accumulation of vesicles in the tube apex is related to the pollen tube growth that depends on active endo/exocytosis (Hála et al., 2008; Hepler and Winship, 2015; Bloch et al., 2016; Marković et al., 2020). The spatial distribution pattern of ANN5 in the elongating pollen tubes indicates that it supports vesicle trafficking, as it was previously reported by Zhu et al. (2014a).

**Figure 4.**
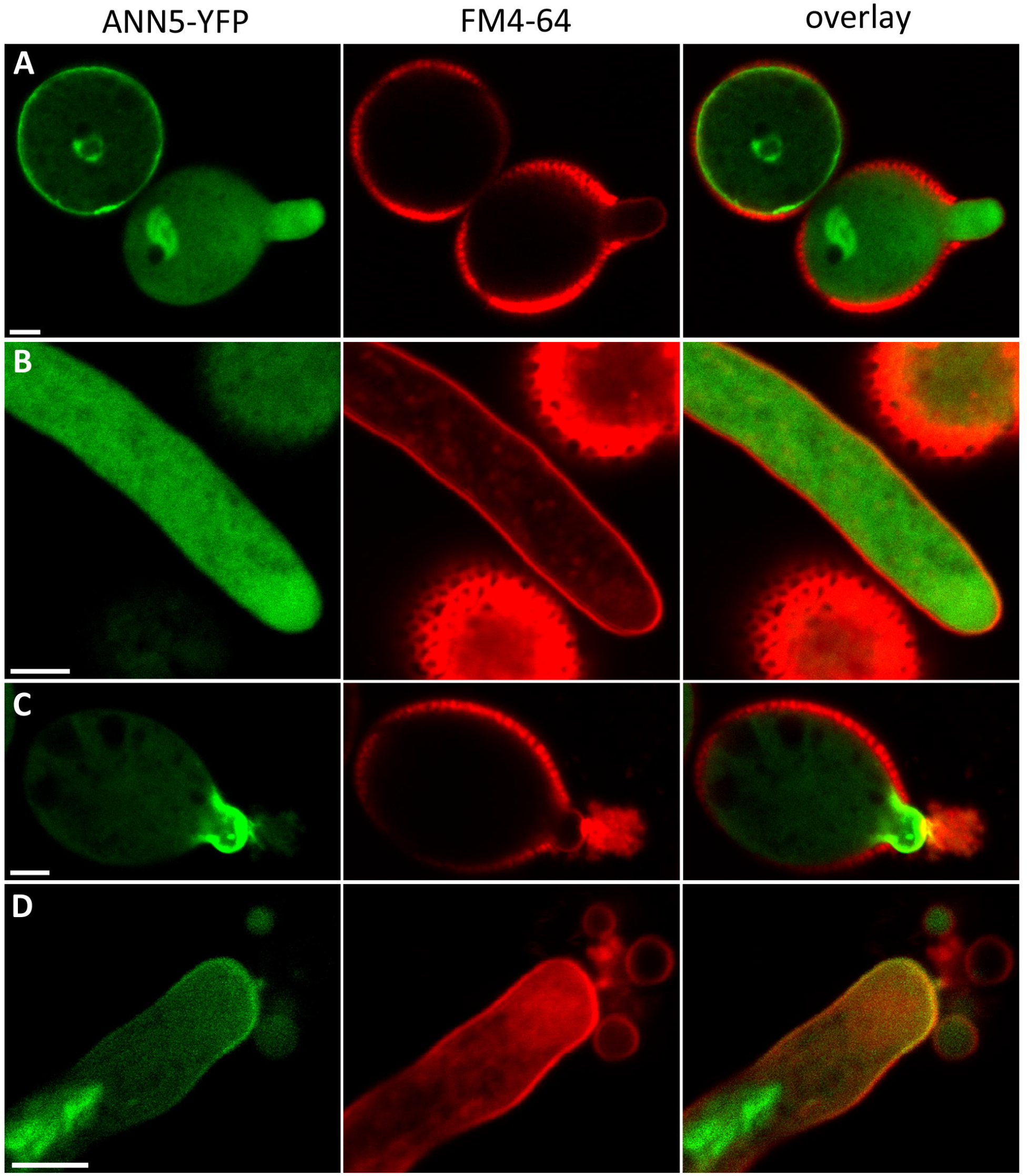
Comparison of ANN5 localization patterns in germinated and non-germinated pollen grains. Representative images of pollen grains overexpressing *ANN5-YFP* after 30 min incubation in liquid pollen germination medium counterstained with FM4-64. (**A**) In osmotically stressed pollen grains ANN5 was associated with PM. In a germinating pollen grain, ANN5 accumulated in the cytoplasm of a newly formed pollen tube. (**B**) In an extending pollen tube ANN5 accumulated in vesicle-rich area at the tube apex. (**C**) A representative confocal image showing pollen tube bursting at the early phase of pollen germination. Formation of a cap-like structure by ANN5 was observed at the PM rupture site. (**D**) Bursting of longer pollen tubes was accompanied by an attachment of ANN5 to the surroundings of the rupture site at the tube apex. Scale bars = 10μm.

We observed that some pollen tubes displayed symptoms of osmotic stress, which was usually accompanied by a bursting of the tube apex. If the pollen tube bursting occurred at the onset of the germination, ANN5 was forming a cap-like structure at the germination site (Fig. 4C). In turn, bursting of the tubes at an advanced stage of the germination resulted in binding of ANN5 to the pollen tube membrane adjacent to the rupture site at the tube apex (Fig. 4D). Although the cytoplasmic leakage was stopped ANN5 remained attached to the pollen tube membrane. Further growth of pollen tube after rupture was not resumed. Massive accumulation of ANN5 at the rupture site and in the surrounding area implicates ANN5 also in the restoration of the pollen tube membrane integrity.

### Ca^2+^ and Mg^2+^ induced pollen membrane defects are accompanied by ANN5 binding

Our findings suggest that PM-association of ANN5 could reflect the osmotically-induced membrane defects. We further tested specific effects of the components of the standard pollen germination medium on ANN5 distribution, because they were shown to affect membrane properties when applied independently (Obermeyer et al., 1996; Pedersen et al., 2006; Schultz et al., 2009; Zhou et al., 2015). Sucrose solutions (20%) supplemented with 1.5 mM Ca(NO_3_)_2_, 1 mM MgSO_4_, 1 mM KNO_3_ or 1.6 mM H_3_BO_3_ were applied to dry pollen grains overexpressing *ANN5-YFP*. To monitor PM permeability, the pollen grains were stained with PI. Interestingly, PI was also shown to enter living cells upon dynamic changes in the membrane potential (Kirchhoff and Cypionka, 2017).

We found that translocation of ANN5 to PM occurred most frequently upon the application of magnesium (Mg^2+^) and calcium (Ca^2+^) ions in comparison to other treatments, i.e., in approx. 70% and 50 % of pollen grains, respectively (Fig. 5). We observed however, opposite effect of these ions on pollen physiology. Exogenously applied Ca^2+^ evoked bursting of pollen grains accompanied by PI influx into the pollen cytoplasm (Fig. 5A). Strikingly, local accumulation of ANN5 at the site of PM damage preceded shortly ejection of the cytoplasmic content of some grains (grain 3, Supplementary Fig. S2). Afterwards ANN5 progressively accumulated at the entire surface of the PM. Other grains collapsed immediately following the bursting (grain 1, Supplementary Fig. S2). In those grains ANN5 was attached to the disrupted PM. There was also a subset of grains in which ANN5 presence on PM was limited to the duration of pollen bursting. Afterwards ANN5 was released rapidly from the membrane possibly as a result of new osmotic equilibrium state that had been reached (grain 2, Supplementary Fig. S2). In contrast, there was no bursting of pollen grains when Mg^2+^ were exogenously applied. However, PM of most pollen grains became permeable to PI that was accompanied by binding of ANN5 to PM (Fig. 5A and 5B). Incubation of pollen grains in sucrose solution containing boric acid (very weak acid) prevented the pollen bursting and consistently the number of grains with ANN5 attached to PM was the lowest in comparison to other treatments. Similarly the addition of potassium ions (K^+^) reduced osmotic-driven pollen rupture and halved the number of grains with PM-associated ANN5 in comparison to the control sucrose solution.

**Figure 5.**
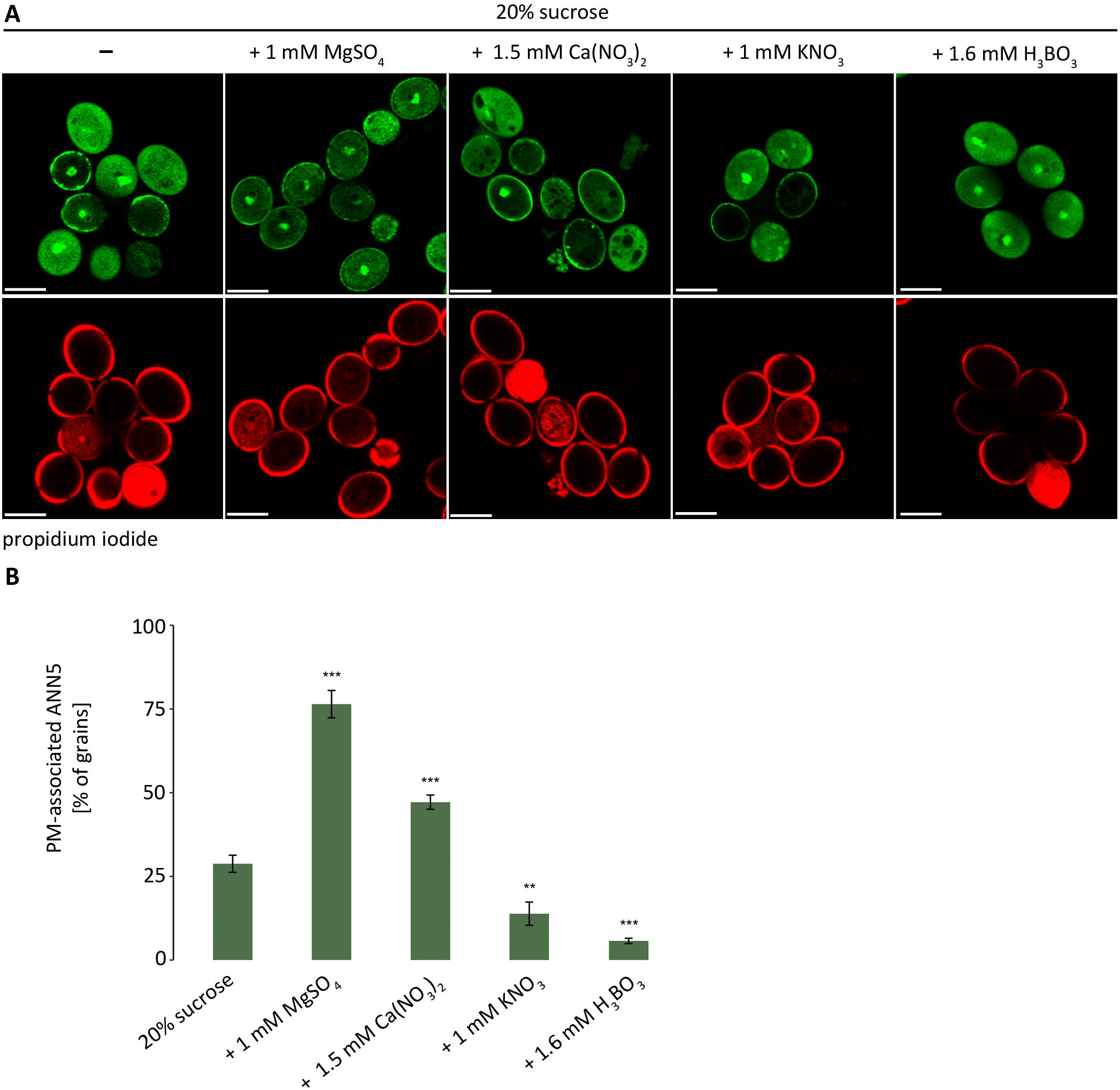
Various effects of components of pollen germination medium on PM permeability and ANN5 distribution. **(A)** Representative images of pollen grains overexpressing *ANN5-YFP* acquired after 30 min incubation in 20% sucrose solutions containing individual compounds and PI. (**B**) Fraction of pollen grains [%] showing PM-associated ANN5 after treatment with different compounds. The data were obtained in three independent experiments, n ≥ 400 pollen grains in each solution. One-way ANOVA with Tukey HSD test was used for statistical analyses. Error bars depict standard deviation. Asterisks indicate means statistically significant, when compared to the control solution. *** p-value < 0.001, ** p-value < 0.01. See also Supplementary Fig. S2.

### Pollen grains dehydrate in response to exogenously applied Mg^2+^

Time-lapse imaging of the pollen grains incubated in the medium containing Mg^2+^ revealed that the number of PM-associated ANN5 grains was increasing with the duration of the experiment (Fig. 6A). However, a 2D-projection of image-stacks showed that the pattern of ANN5 distribution within PM varied depending on the external conditions (Fig. 6C). In pollen grains incubated in sucrose solution numerous punctuate signals were detected in addition to diffuse fluorescence of ANN5-YFP in the plane of pollen membrane, suggesting ANN5 presence in PM subcompartments. After exposure to Mg^2+^ ions the initial diffuse distribution of ANN5 at PM changed quickly into more patchy pattern as a result of ANN5 clustering at PM. Moreover, association of ANN5 with PM correlated with progressive shrinkage of the pollen grain volumes in response to. We wondered whether the decrease in pollen grain volumes induced by Mg^2+^ might be reminiscent of that usually observed under salt stress. Therefore we examined an effect of 150 mM NaCl on pollen hydration status and ANN5 distribution. Shrinkage of pollen grains was observed after 35 min of incubation in 150 mM NaCl (by approximately 7%, Supplementary Fig. S3C) which is comparable to that induced by Mg^2+^ (ca. 9%, Fig. 6B). However, the association of ANN5 with PM was scarce under salt stress. ANN5 accumulated primarily in the cytoplasm in close vicinity to the PM area permeable to PI (Supplementary Fig. S3A). Next, ANN5 was redistributed non-uniformly within the vegetative cell and organized into an interconnected mesh-like structure (Supplementary Fig. S3A, S3B), suggesting that pattern of ANN5 distribution in response to Mg^2+^ is specific.

**Figure 6.**
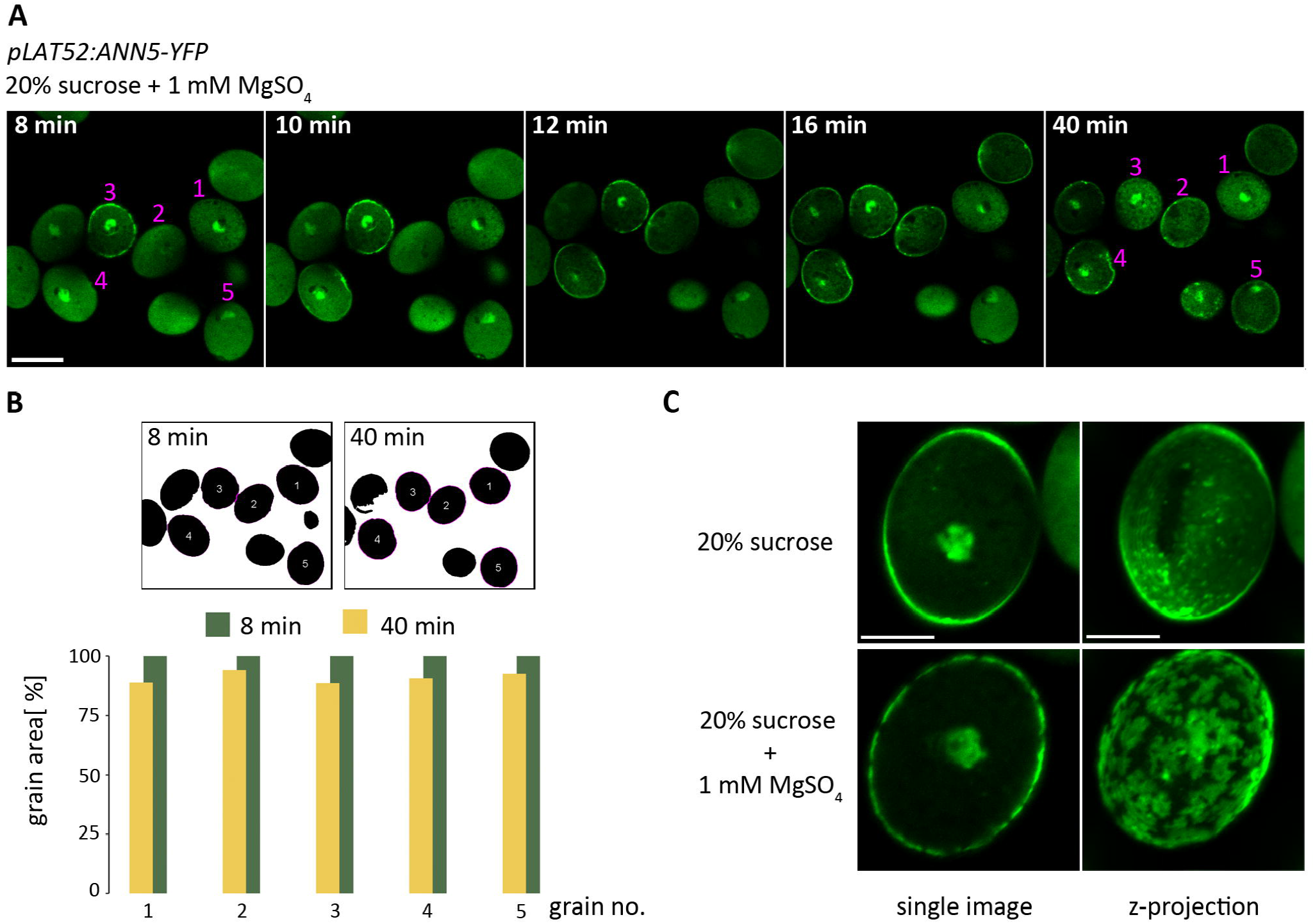
Pollen grains dehydration in response to external Mg^2+^ is accompanied by binding of ANN5 to PM. **(A)** Representative time-lapse images of pollen grains overexpressing *ANN5-YFP* acquired during incubation in 20% sucrose solution with the addition of 1 mM MgSO_4_. A scale bar = 20 μm. (**B**) A grain area was measured for 5 individual pollen grains (depicted with numbers) on single confocal images acquired at 8 and 40 min time points. The normalized values plotted on the graph show a reduction in pollen grain area by ca. 9%. Compare with dehydrating effect of 150 mM NaCl provided in Supplementary Fig. S3. **(C)** Clustering of ANN5 on PM after 40 min incubation in the presence of external Mg^2+^. Representative images and z-projections of pollen grains incubated in the control solution (20% sucrose) and with the addition of Mg^2+^. Z-projection is 2D maximum intensity projection of z-stack consisting of 13 images generated in ImageJ. Scale bars = 10 μm.

To check whether Mg^2+^-induced PM permeability is a general phenomenon, we analyzed PI influx in pollen of *Brasica napus*, another species from the *Brassicaceae* family, as well as pollen of *Nicotiana tabacum*, a representative from *Solanaceae* family and *Malus domestica*, a representative from *Rosaceae* family during exposure to the exogenous Mg^2+^. As shown in Supplementary Fig. S4, the response of *B. nopus* pollen grains was similar to Arabidopsis, that is Mg^2+^-induced membrane permeabilization and cellular dehydration observed as PI influx and a reduced grain size, respectively (Supplementary Fig. S4B). In contrast, we did not detect such changes in the pollen grains derived from *N. tabacum* and *M. domestica* exposed to Mg^2+^ (Supplementary Fig. S4C).

### Mg^2+^ induce pollen membrane depolarization

In order to characterize the effect of individual components of the standard pollen germination on PM properties, we used two high voltage-sensitive dyes to track changes in the membrane potential. Cationic DiOC(2)3 enters a cell due to its affinity for the negatively polarized membranes, and accumulates in the energized and negatively charged mitochondria. The anionic DiBAC4(3) is excluded by a polarized cell, because it is negatively charged and its influx into the cell indicates depolarization of the membrane. The staining was performed only on the Arabidopsis Col-0 pollen grains because the fluorescence emission spectra of the dyes DiOC(2)3 and DiBAC4(3) overlap with YFP emission spectrum.

We observed that the PM resting potential was not affected by the incubation in the control sucrose solution or in the presence of 1.5 mM Ca^2+^ since DiOC(2)3 entered the majority of the pollen grains and accumulated in the mitochondria, whereas DiBAC4(3) remained outside the grains (Fig. 7A, 7B). However, there was always a subset of the grains that were permeable to DiBAC4(3) but DiOC(2)3 was not accumulating in the mitochondria. Thus we conclude that in those grains osmotic stress elicited PM defects. In contrast, the majority of pollen grains treated with Mg^2+^ displayed depolarization of PM visualized by an influx of the DiBAC4(3) and inhibition of DiOC(2)3 accumulation in mitochondria. In turn, in the presence of boric acid or K^+^ we observed reduction of DiOC(2)3 influx and retention of DiBAC4(3) outside of the pollen grains indicating PM hyperpolarization. In the standard pollen germination medium pollen grains germinate and form pollen tubes indicating that the adverse effects of individual compounds are eliminated when used in a combination.

**Figure 7.**
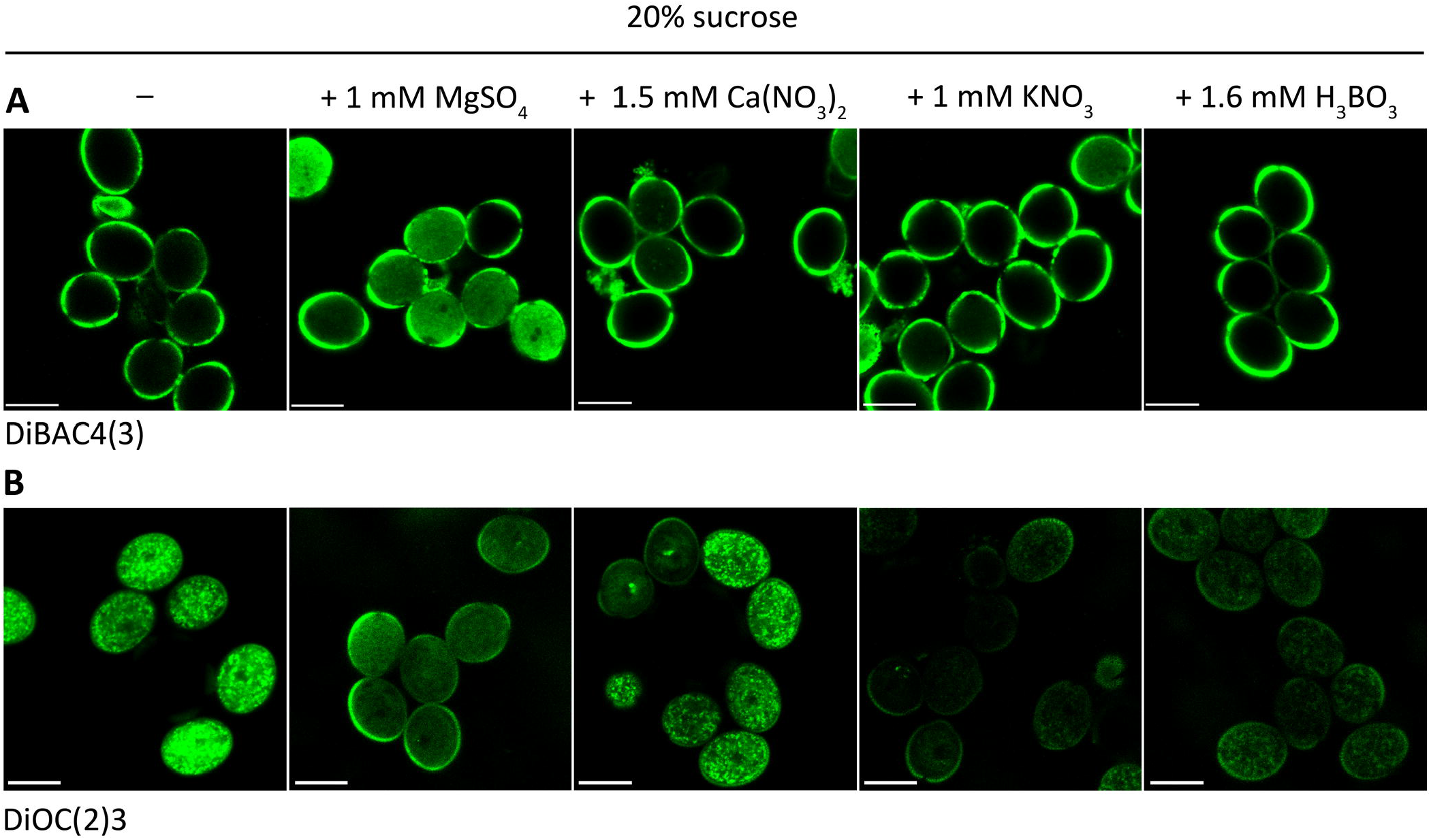
The effects of components of pollen germination medium on PM properties. Representative images of Arabidopsis (Col-0) wild-type pollen grains acquired after 30 min incubation in 20% sucrose solutions containing different compounds, counterstained with fluorescent dyes: DiBAC4(3) **(A)** or DiOC(3)2 **(B)**. Scale bars = 20 μm.

### The effects of Mg^2+^ on pollen grains are eliminated in the presence Ca^2+^ and K^+^

In order to find out which component of the pollen germination medium antagonizes the effects of Mg^2+^, pollen grains overexpressing *ANN5-YFP* were incubated in the sucrose solution containing Mg^2+^ and one of the other compounds. Time-lapse imaging was started 10 min after immersion of the pollen grains in the appropriate solution. Confocal images were acquired every 1 min over a total of 55 min for each experiment. Counterstaining with PI was included for tracking the kinetics of PM permeabilization during incubation of the pollen grains in the solutions tested. In the Mg^2+^-containing sucrose solution the accumulation of ANN5 at PM preceded shortly PM permeability for PI (Fig. 8C). Once the clusters of ANN5 at PM were formed we observed a progressive influx of PI into the pollen cytoplasm. Over time the pollen grains reduced their size due to the water loss whereas ANN5 remained attached to the PM (Fig. 8A and 8E), as we have observed earlier (Fig. 6).

**Figure 8.**
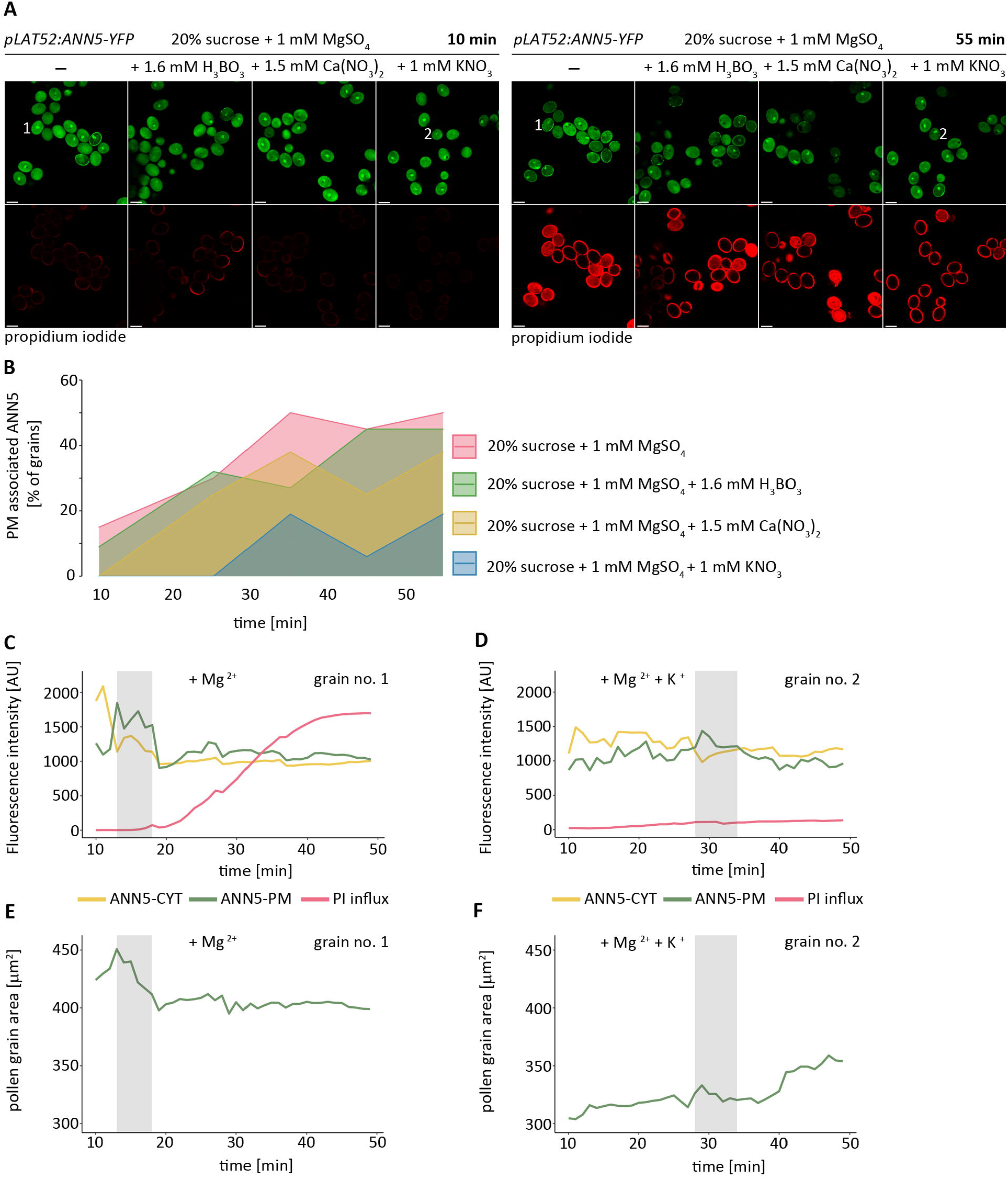
The antagonistic effects of Mg^2+^ and K^+^ in mature pollen grains. **(A)** Representative images of pollen grains overexpressing *ANN5-YFP* after 10 and 55 min incubation in sucrose solutions containing 1 mM MgSO_4_ or supplemented with the additional compound and stained with PI. See also Supplementary Video S2. Pollen grains marked with numbers (1 and 2) were selected for the measurement of the relative fluorescence intensities presented on graphs in sections C, D, E and F. Scale bars = 20 μm. **(B)** Graph showing the change in the number of PM-associated ANN5 pollen grains [%] during incubation in sucrose solution containing Mg^2+^ or supplemented with the additional compound. Data from one representative experiment of three independent experiments is shown, n = 16-22 pollen grains. Plots of the change in the relative fluorescence intensities of ANN5-YFP and PI measured in pollen cytoplasm (ANN5-CYT and PI) and pollen membrane (ANN5-PM) during incubation in sucrose solution containing Mg^2+^ **(C)** or Mg^2+^ and K^+^ **(D).** Plots of the change in the pollen grain area [μm^2^] during incubation in sucrose solution containing Mg^2+^ **(E)** or Mg^2+^ and K^+^ **(F)**. The gray shaded region on the graphs (C, D, E, and F) indicates time of ANN5 accumulation at PM [min].

The addition of boric acid did not eliminate the impact of Mg^2+^ (Fig. 8A and 8B, Supplementary Video S2). The cellular dynamics of ANN5 as well as influx of PI were like in the Mg^2+^-containing sucrose solution. In contrast, the addition of Ca^2+^ eliminated the effects of Mg^2+^. This was, however, accompanied by an increased number of the pollen bursting events resulting in PM rupture, PI influx and translocation of ANN5 to the PM damage site. Time of ANN5 attachment to PM in those pollen grains was limited to the duration of pollen bursting. Such effects were commonly observed in the pollen grains incubated in the sucrose solution containing solely Ca^2+^. The addition of K^+^ abolished the negative effects of Mg^2+^in the majority of pollen grains. K^+^ ions positively influenced PM integrity by preventing PI influx and reducing the number of PM-associated ANN5 pollen grains (Fig. 8A). Only a small number of the pollen grains were affected during incubation in the sucrose solution containing Mg^2+^ and K^+^, identified by a delayed occurrence of ANN5 at PM. There was also a small fraction of the pollen grains showing transient association of ANN5 with PM. Such pollen grains did not dehydrate, however, a slight PI influx was detected over time of livecell imaging (Fig. 8D). After dissociation of ANN5 from PM we observed an increase in the pollen grain size (Fig. 8F). This finding suggests that pollen grains were initially affected by Mg^2+^ however this effect was reversed due to the presence of K^+^ (Supplementary Video S2).

### Pollen developmental stage-dependent sensitivity to Mg^2+^

Our data indicate that Mg^2+^ affect mature pollen grains in Arabidopsis and *B. napus*. To determine if Mg^2+^ exerts negative effects also on pollen grains before anthesis we performed analogous experiments on pollen grains collected from the final bud stage 12, according to the classification described previously (Sanders et al., 1999). The pollen grains of the wildtype and overexpressing *ANN5-YFP* lines were incubated in 300 mM mannitol containing 1 or 5 mM Mg^2+^ for 30 min. Counterstaining with PI allowed us to simultaneously visualize PM permeability. Mg^2+^ induced PM permeability in both wild-type and transgenic pollen grains collected from dehisced anthers as seen by an influx of PI into the cytoplasm and a translocation of ANN5 to PM (Fig. 9B). Surprisingly the integrity of PM was not affected even in the presence of 5 mM Mg^2+^ in the pollen collected from the flower buds, derived from wild-type as well as transgenic Arabidopsis plants (Fig. 9A). Neither PI influx nor ANN5 translocation in response to the elevated Mg^2+^ levels were observed indicating that sensitivity of PM to Mg^2+^ depends on the developmental stage of the pollen grain.

**Figure 9.**
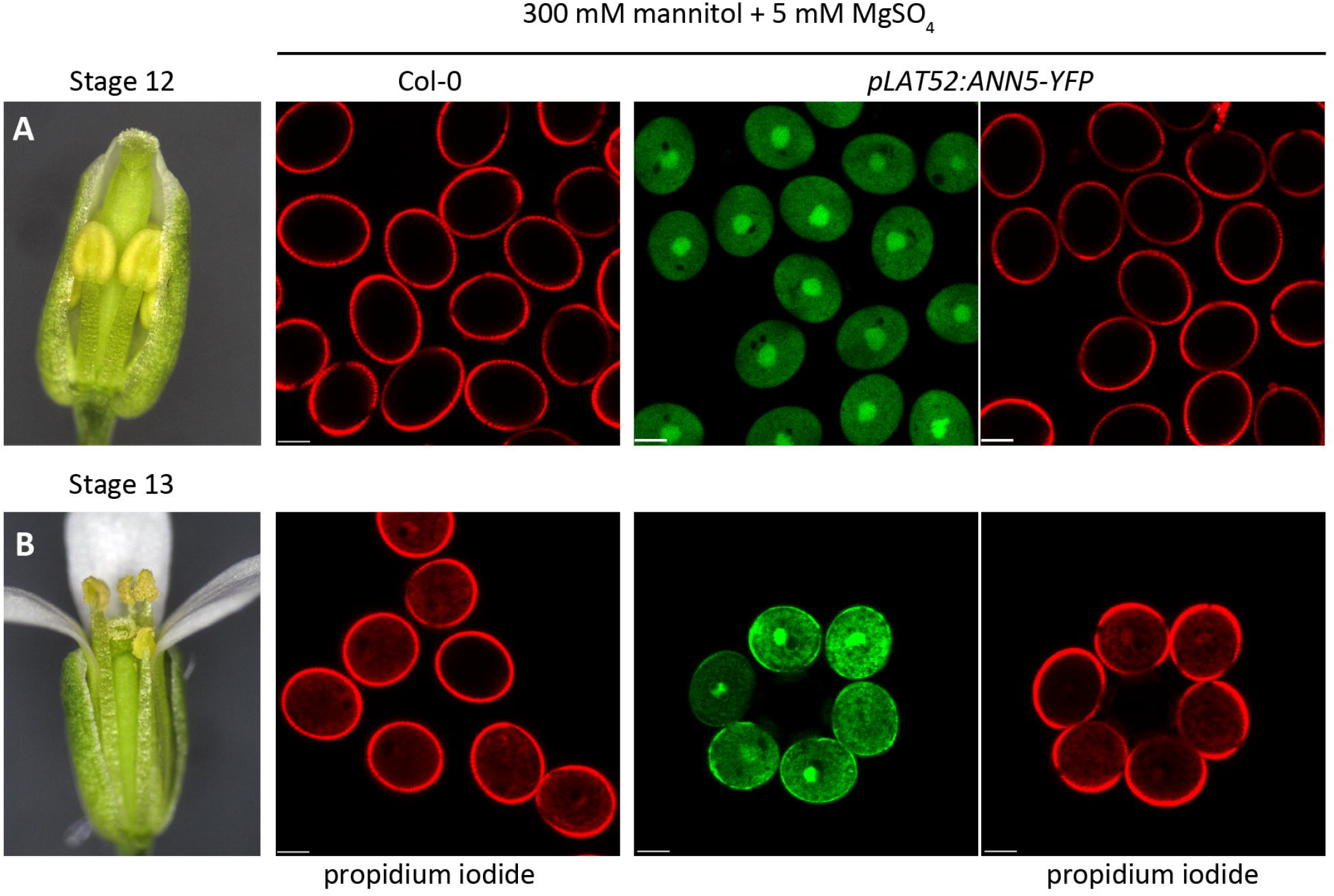
Analysis of male gametophyte developmental stage-dependent sensitivity to magnesium. Representative images of Arabidopsis pollen grains of wild-type (Col-0) and overexpressing *ANN5-YFP* incubated in 300 mM mannitol solution with addition of 5 mM MgSO_4_ for 30 min. Scale bars = 10 μm. (B) Mg^2+^ induced membrane permeability for PI in mature pollen grain collected from open flowers (stage 13). Influx of PI was accompanied by ANN5 binding to the PM. See also response of *B. napus* mature pollen grains to exogenous Mg^2+^ (Supplementary Fig. S4). (A) Mg^2+^ did not affect PM integrity in pollen grains from the late flower bud (stage 12) of wild-type and overexpressing *ANN5-YFP*, observed as a lack of PI staining. Compare with PI influx in lines silenced for *ANN5* in Supplementary Fig. S5.

Collectively, we showed developmentally regulated change in the receptivity of the pollen grains to exogenous Mg^2+^.

We showed that pollen grains from the late flower buds were not receptive to Mg^2+^. We wondered whether *ANN5* expression levels may determine pollen sensitivity to exogenous Mg^2+^. Therefore we analyzed PI influx in the presence or absence of 5 mM Mg^2+^ in the pollen grains isolated from the late flower buds of the wild-type Arabidopsis, *ANN5* RNAi-silenced and overexpressing *ANN5-YFP* lines, that were generated in our previous (Lichocka et al., 2018) and this study, respectively. The fraction of PI-stained pollen grains was much higher in the lines silenced for *ANN5* in comparison to the wild-type or overexpressing *ANN5-YFP* lines exposed to Mg^2+^. This might suggest that ANN5 provides protection to exogenous Mg^2+^ in the pollen grains from the late flower buds (Supplementary Fig. S5). However the number of PI-stained pollen grains in *ANN5* RNAi-silenced lines was also higher in the control sucrose solution comparing to the wild-type. This observation is consistent with the results of our previous studies showing abnormalities in the generative development in *ANN5* silenced lines (Lichocka et al., 2018).

## Discussion

The involvement of plant annexins in the maintenance of plasma membrane integrity has long remained largely elusive. By contrast, the important role of annexins in various steps of the membrane repair process is well demonstrated in mammalian cells in a number of research papers. Annexins directly strengthen the membrane at an injury site as well as orchestrate assembly of other proteins involved in removal of the membrane wound (Creutz et al., 2012; Boye and Nylandsted, 2016; Demonbreun et al., 2019; Sonder et al., 2019). A self-organization of annexins into homo- and hetero-multimers forming a 2D molecular lattice at the membrane surface seems to initiate this process (Zaks and Creutz, 1991; Lin et al., 2020). An ability of some plant annexins to oligomerize *in vitro* suggests that they may function like their metazoan counterparts (Hoshino et al., 1995; Hofmann et al., 2002; Gorecka et al., 2005; Konopka-Postupolska et al., 2009) assembling into higher ordered structures at membranes. Importantly, the data presented in this study demonstrate that either osmotic or ionic stress may trigger the recruitment of ANN5 to the membrane of pollen grains in Arabidopsis. Our findings strongly support the concept that at least some plant annexins play a role in the membrane repair.

In this study, we focused on the role of ANN5, a pollen-specific annexin, during *in vitro* and *in vivo* hydration and germination of pollen grains. Live-cell imaging experiments revealed that ANN5 associates with PM during a rapid increase in the pollen grain volume that results in PM overstretching. Such effects were induced in the pollen grains during *in vitro* hydration and after spraying on the pollinated stigma with deionized water (Fig. 1, Fig. 2 and Fig. 3). Rapid hydration of the pollen grains similarly to seeds imbibition may change plasma membrane permeability and provoke leakage of the solutes (Simon and Mills, 1983; Crowe et al., 1989; Golovina et al., 1998). Based on our findings, we propose that ANN5 strengthens PM exposed to overstretching during sudden change in water balance that under natural conditions might be induced by rainfalls.

A prolonged osmotic stress led to the loss of membrane integrity manifested by bursting of pollen grains and pollen tubes (Fig. 4). Our data show that ANN5 massively accumulated at the membrane rupture sites. This pattern of ANN5 distribution resembles a repair cap structure, a macromolecular repair complex that is formed by the metazoan annexins at the membrane damage sites (Zilly et al., 2011; Demonbreun et al., 2016; Demonbreun et al., 2019). This structure serves as an assembly platform for other proteins assisting in the membrane repair (Sonder et al., 2019). We propose that formation of the cap-like structure by ANN5 and binding to the membrane surrounding the rupture site are two necessary steps of rapid restoration of the membrane integrity in pollen grains and pollen tubes.

We also show that ANN5 is involved in cellular response to ionic stress exerted by externally applied Mg^2+^ or NaCl (Fig. 5, Fig. 6, Fig. 7, Fig. 8, Supplementary Fig. S3). Both treatments enhanced PM permeability and induced pollen grains dehydration but differently affected the cellular distribution of ANN5. Mg^2+^ induced binding of ANN5 to PM whereas NaCl forced ANN5 association with the intracellular structures. We thus infer that ANN5 distribution patterns reflect differences in the cellular response to varying extracellular conditions. An important role of plant annexins in conferring stress tolerance was previously reported (Szalonek et al., 2015; Zhang et al., 2015; Ijaz et al., 2017), however, their stimulus-dependent cellular relocation has not been demonstrated yet. In contrast, it is well established that metazoan annexins in response to stress are mobilized in a calcium-dependent manner to the plasma membrane, (Boye and Nylandsted, 2016; Boye et al., 2017; Demonbreun et al., 2019; Sonder et al., 2019; Lin et al., 2020). There is also evidence of a nuclear translocation of human ANXA2, in a phosphorylation-dependent manner, under heat and oxidative stress conditions (Rhee et al., 2000; Liu et al., 2003; Grindheim et al., 2017). These findings indicate that annexins may serve as stress-responsive proteins providing a cellular tolerance to adverse conditions. Based on our results we propose that ANN5 is implicated in the maintenance of cellular homeostasis in the mature pollen grains. However further studies are required to elucidate the underlying mechanisms.

We showed that exogenously applied Mg^2+^ induced dehydration of pollen grains not only in Arabidopsis but also in *B. napus* plants (Fig. 6, Fig. 8 and Supplementary Fig. S4). Arabidopsis and *B. napus* represent a model of dry stigma that in response to pollination provides water to pollen grains in a controlled manner. Interestingly, treatment with Mg^21^ did not affect pollen grains from plants representing wet-stigma type such as *N. tabacum* and *M. domestica* (Supplementary Fig. S4) in which after pollination the pollen grains acquire ions from the pistil exudates. We therefore propose that in *Brassicaceae*, pistil-derived Mg^2+^ might be another important player involved in pollen-pistil interaction, potentially regulating hydration level of the pollen grains on stigma. A growing body of evidence indicates that ionic components (Ca^2+^ and K^+^) derived from stigma are key regulatory factors during pollen-pistil interaction and may contribute to self-pollen rejection (Iwano et al., 2015; Ma et al., 2018). Our findings reveal another level of complexity in pollen-pistil interaction and open new exciting avenues for exploring a role of ion dynamics in the pollen recognition.

In our study the Mg^2+^ impact on pollen grains was eliminated in the presence of Ca^2+^ and K^+^ (Fig. 8). Similar phenomenon was previously reported in *Crinum asiaticum* (Kwack, 1964) showing that inhibitory effect of Mg^2+^ on pollen germination could be eliminated by the addition of Ca^2+^. Our result is a another example of the antagonism between Ca^2+^ and Mg^2+^during pollen germination *in vitro* and it is in agreement with the current knowledge showing that the intracellular homeostasis depends on Ca^2+^/ Mg^2+^ equilibrium (Tang and Luan, 2017). The competition between Mg^2+^ and K^+^ was mostly studied in plant roots and leaves (Xie et al., 2020). It was shown that high K^+^ to Mg^2+^ ratio interferes with Mg^2+^ uptake whereas high level of Mg^2+^ increases the risk of K^+^ deficiency (Osterhout, 1908; Gransee and Führs, 2013; Laekemariam et al., 2018). Our data suggest that similar relationship between Mg^2+^ and K^+^ may occur in pollen grains.

So far it was shown that disruption of specific Mg cations transporters in Arabidopsis *(Atmgt5, Atmgt9* and *Atmgt4)* led to pollen abortion (Li et al., 2008; Chen et al., 2009; Li et al., 2015) suggesting an important role of Mg^2+^ during pollen development (Mueller-Roeber and Arvidsson, 2009). In our study application of Mg^2+^ did not affect the pollen grains isolated from flower buds whereas in the mature pollen grains led not only to PM depolarization but also induced shrinkage of the PM surface area as a result of water loss (Fig. 6, Fig. 7 and Fig. 9). Further research is needed to understand a detailed role of Mg^2+^ at different stages of pollen development. Here, we show that under stress conditions ANN5 displayed PM localization (Fig. 6). The initial diffuse distribution of ANN5 gradually changed into distinct punctuate subdomains or larger clusters during incubation in the sucrose solution or in the presence of Mg^2+^ ions, respectively. Segregation of specific lipids along with the assisted proteins into membrane distinct domains usually takes place during signal transduction and membrane trafficking events (Jaillais and Ott, 2020). It was previously shown that membrane depolarization also led to clustering of the negatively charged phospholipids such as phosphatidylserine accompanied by co-clustering of the membrane-related proteins (Zhou et al., 2015). Cell shrinkage also drove structural reorganization of the membrane by forcing higher lipid packing density (Janmey and Kinnunen, 2006). Recent studies shed light on the importance of spatial reorganization of membrane lipids during development of pollen grains and growth of pollen tubes (Lee et al., 2018; Fratini et al., 2020; Zhou and Dobritsa, 2020). Annexins bind preferentially anionic phospholipids and some members have been shown to associate with the membrane subdomains (Demir et al., 2013; Drücker et al., 2013). We therefore propose that the formation of distinct subdomains containing ANN5 within PM reflects lateral reorganization of the anionic phospholipids in the inner leaflet of PM. This mechanism would account for the possible ANN5 function in Ca^2+^ signaling and intracellular membrane trafficking in mature pollen grains (Ruan et al., 2021).

Taken together, our experiments indicate that manipulation of osmotic and ionic balance may induce mechanical tension in PM that may finally result in its local damage. We showed the ability of ANN5 to associate with PM, which integrity is compromised or destroyed. This phenomenon has not been previously described for other members of the plant annexin family. Further analysis will be required to establish physiological consequences of ANN5 assembly at the plasma membrane and to verify its putative involvement in the membrane repair processes. Our study provides also a tool to track PM damage *via* live-cell imaging of ANN5 which may be employed to gain a better insight into the mechanisms of membrane repair in plant cells.

## Data Availability

GenBank accession numbers of gene sequences used in this study: ANN5 (AT1G68090), ANN1 (AT1G35720). The datasets generated for this study are available on request to the corresponding author.

## Author Contributions

M.L. designed the study, performed research and wrote draft manuscript. M.K. edited the manuscript and gave conceptual advices. M.G. developed charts and conducted all statistical analyses. J.H. supervised the study and gave conceptual advices.

## Conflict of Interest

The authors declare that the research was conducted in the absence of any commercial or financial relationships that could be construed as a potential conflict of interest.

## Supplementary Materials

**Supplementary Figure S1.** Uptake of PI occurs in pollen gains with ANN5 attached to PM.

**Supplementary Figure S2.** Ca^2+^ application induces pollen bursting accompanied by ANN5 binding to PM.

**Supplementary Figure S3.** Pollen grains dehydration upon exposure to NaCl.

**Supplementary Figure S4.** Analysis of PI uptake by *B. napus, N. tabacum* and *M. domestica* mature pollen grains upon exposure to exogenous Mg^2+^.

**Supplementary Figure S5.** Analysis of PI uptake in pollen grains from flower buds (stage 12) in Arabidopsis with modified *ANN5* expression levels.

**Supplementary Video S1.** Translocation of ANN5 to PM in pollen grains germinating on stigma after placing a drop of deionized water.

**Supplementary Video S2.** ANN5 dynamics in pollen grains incubated in Mg^2+^-containing sucrose solution or supplemented with putative Mg^2+^ antagonizers.

## Notes

### Competing Interest Statement

The authors have declared no competing interest.

